# PHYRE, a Translated uORF, is Required for Phytochrome B Translation and Photomorphogenesis in Arabidopsis

**DOI:** 10.64898/2026.10.31.763305

**Authors:** Naveen Shankar, Anne Van Humbeeck, Manuela Jurca, Ashleigh Edwards, Amir Mahboubi, Johannes Hanson, José Carlos Montanés, Chris Papadopoulos, M. Mar Albà, Stephan Wenkel

## Abstract

Translational control allows cells to adjust protein output independently of transcription, yet the mRNA-encoded mechanisms that govern this regulatory layer remain poorly understood. Upstream open reading frames (uORFs) are among the most prevalent translational regulatory elements in eukaryotic genomes; however, characterised examples of activating uORFs that promote rather than inhibit downstream translation are exceedingly rare. Here we show that efficient expression of phytochrome B (phyB), the master red-light photoreceptor of Arabidopsis thaliana, requires translation of a short uORF in its own 5′UTR, which we name *PHYTOCHROME REGULATOR* (*PHYRE*). Loss of *PHYRE* function phenocopies *phyB* null mutants across germination, hypocotyl growth, leaf development, and flowering time. *PHYRE* activity depends on its encoded peptide sequence, and the spacing between *PHYRE* and the *PHYB* start codon is a critical determinant of downstream translation efficiency, consistent with a position-dependent reinitiation mechanism. *PHYRE*-like elements are conserved across eudicots, establishing uORF-mediated translational gating as a broadly maintained regulatory layer controlling a master photoreceptor.

## INTRODUCTION

Eukaryotic cells can respond rapidly to signals, and within seconds strong changes in the transcriptome can be observed that rely predominantly on transcription^1^. However, mRNA abundance and protein output are frequently decoupled, and transcripts that accumulate rapidly are not necessarily translated. Systematic comparisons of mRNA and protein levels across eukaryotes have established that transcript abundance explains only approximately 40-50% of the variation in protein levels^2,3^. The remainder reflects additional regulatory processes, including translational control, post-translational modifications, and protein degradation, that collectively shape protein abundance and activity but remain less understood. Ribosome profiling, which captures the positions of translating ribosomes at genome-wide scale, has revealed extensive translational regulation that is uncoupled from changes in mRNA abundance^4,5^. Mechanistic models further demonstrate how translation initiation, elongation and ribosome traffic jointly determine protein synthesis rates^6,7^. Moreover, translational activity is dynamically reprogrammed in response to environmental cues and temporal changes in plant physiology^8,9^. Translational control thus constitutes an independent regulatory axis through which cells modulate their protein output, and understanding its molecular basis remains a central challenge in cell and developmental biology.

One mechanism by which this regulation is encoded directly within mRNA sequences involves upstream open reading frames (uORFs) located in the 5′ untranslated region (UTR). During ribosome scanning, initiation at an uORF can cause the ribosome to stall, terminate, or reinitiate at a downstream position, thereby modulating translation of the primary coding sequence^10^. uORFs are prevalent across eukaryotic genomes, occurring in the 5′UTRs of 30-50% of mammalian and plant transcripts^11,12^. One of the best-studied examples of uORF-mediated translational regulation is of *GCN4*, where (uORFs) inhibit translation under non-stress conditions but allow increased *GCN4* translation during amino acid starvation^13^. Inhibitory uORFs that operate through nascent peptide-mediated ribosome stalling have also been described^14^. By contrast, uORFs that directly activate translation of a downstream ORF are exceedingly rare, and no clear mechanistic paradigm for an activating uORF has been established in plants or animals^12,15^. Consequently, uORF-encoded peptides as a class of functional regulators that promote rather than inhibit downstream expression remain almost entirely uncharacterised^16^.

Plants must continuously monitor their light environment and adjust development accordingly, as light is both their primary energy source and a key positional signal. Among the photoreceptors that mediate light perception, phytochrome B (phyB) functions as a master developmental switch. The phyB protein is activated by red light, which converts it to its active Pfr form, and inactivated by far-red light, reverting phyB to its inactive Pr form^17,18^. Under conditions in which the ratio of red to far-red light is reduced, as occurs when plants grow close to competitors or beneath a plant canopy, active phyB levels fall and trigger the shade avoidance syndrome, a coordinated developmental programme that includes hypocotyl elongation, altered leaf morphology, and accelerated flowering^19^. These diverse developmental outputs place phyB protein abundance at the centre of multiple critical life history decisions. Notably, *PHYB* mRNA is constitutively expressed and does not vary markedly with light conditions, yet phyB protein abundance is tightly regulated across developmental stages and in response to the light environment^20,21^. This discordance between stable transcript levels and regulated protein output suggests that post-transcriptional mechanisms contribute substantially to governing phyB expression.

Despite decades of investigation, the mechanisms controlling phyB protein levels have been studied almost exclusively at the transcriptional and post-translational levels. At the post-translational level, phyB abundance and activity are regulated through light-dependent nuclear accumulation, interactions with PHYTOCHROME INTERACTING FACTOR (PIF) transcription factors, and ubiquitin-mediated degradation^21,22,23^. Whether translational control contributes to the regulation of phyB protein expression has not been examined. Ribosome profiling has recently been applied in plants and revealed a complex translational landscape that includes numerous small ORFs in 5′UTRs without known function^8,24,25^. Whether any of these novel translated elements regulate the expression of major developmental regulators has remained an open question.

Here, we identify a translated upstream ORF in the *PHYB* 5′UTR, which we name *PHYTOCHROME REGULATOR* (*PHYRE*), through ribosome profiling of Arabidopsis seedlings subjected to shade. Unlike the predominant class of inhibitory uORFs, *PHYRE* translation is required to promote phyB protein expression. Loss of *PHYRE* function phenocopies loss of phyB across multiple developmental contexts, including seed germination, hypocotyl elongation, leaf development, and flowering time.

Genetic complementation and reporter analyses demonstrate that *PHYRE* activity depends on its encoded peptide sequence and that the spacing between the *PHYRE* stop codon and the *PHYB* start codon is a critical determinant of downstream translation efficiency. *PHYRE*-like elements are conserved across angiosperms, suggesting that translational activation of *PHYB* by an upstream ORF is a broadly conserved regulatory mechanism. Together, these findings identify *PHYRE* as one of very few characterised activating uORFs in any eukaryote and establish uORF-mediated translational gating as a new regulatory layer controlling a master photoreceptor.

## RESULTS

Despite their prevalence across eukaryotic genomes, short open reading frames (sORFs) remain a largely unexplored layer of translational regulation. To systematically identify translated sORFs in Arabidopsis seedlings exposed to light and shade, we performed ribosome profiling on 7-day-old seedlings grown under white light and subjected to 45 or 90 min of deep shade (Fig. 1A; see Methods). Integrating all nine Ribo-seq datasets, we predicted translated non-canonical sORFs using RibORF v.2.0^26^ (Figs. 1B, S1; see also Methods). This analysis identified 2,856 novel translated sORFs, comprising 2,393 uORFs, 366 ncRNA-associated ORFs, and 97 downstream ORFs (Fig. 1C, Supplementary Table 1). Filtering for high confidence candidates with greater than 50% read periodicity reduced this set to 208 sORFs, including 184 uORFs, 21 ncRNA ORFs, and 3 dORFs, which were prioritised for further analysis (Fig. 1C, Supplementary Table 2).

**Figure 1.**
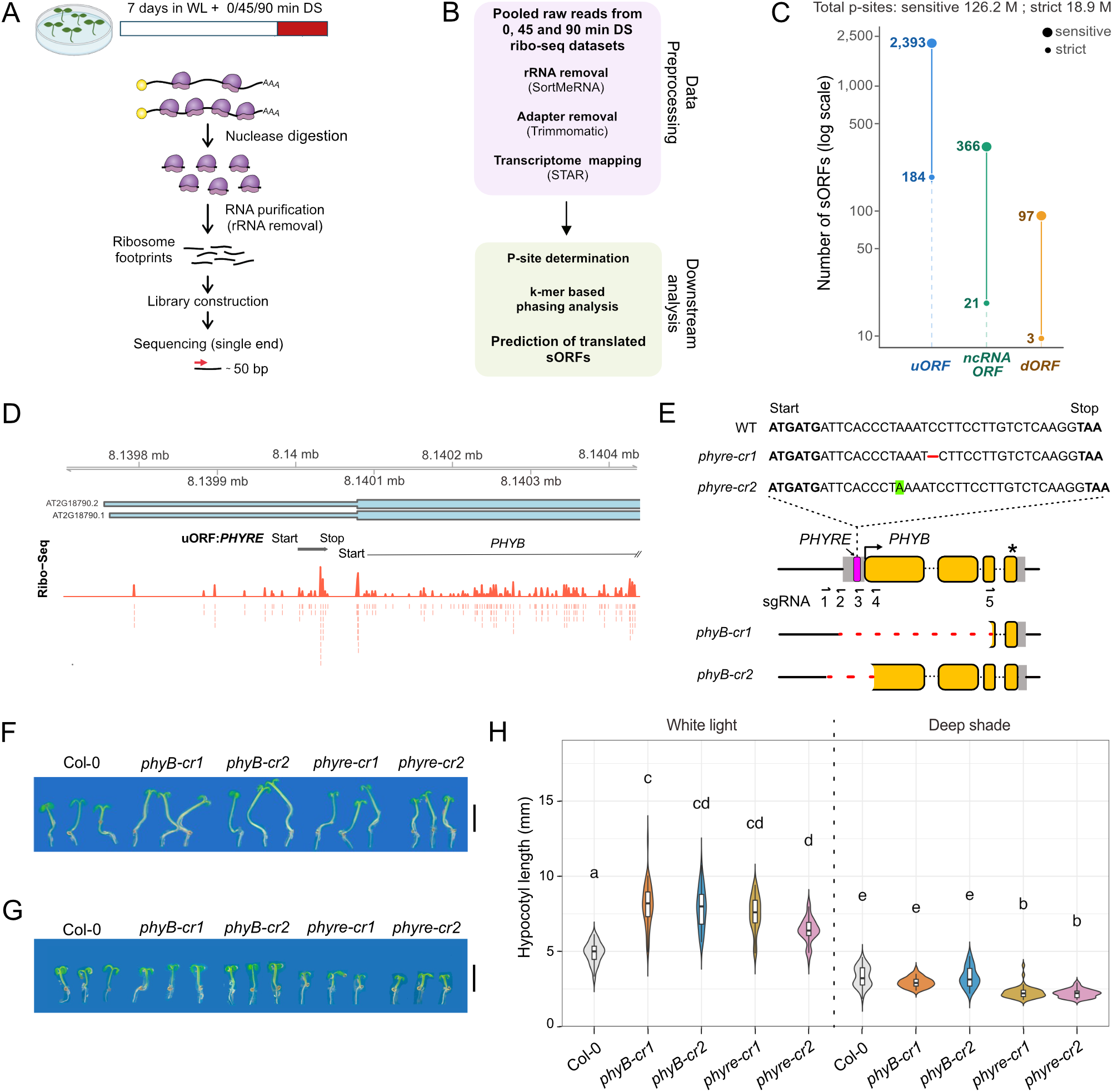
Ribo-seq analysis and hypocotyl phenotypes of *phyB* and *phyre* mutants. **(A)** Schematic of the Ribo-seq workflow. WL = white light, DS = deep shade. **(B)** Schematic of the data analysis pipeline used to discover novel small ORFs. **(C)** Lollipop plot showing the numbers of novel uORFs, non-coding RNA ORFs, and dORFs identified using sensitive and strict analyses. The total number of p-sites determined by sensitive and strict analysis are indicated above the plot. **(D)** Ribosome profile of *PHYRE* and start of the *PHYB* coding region. Transcript gene models in blue are shown below the chromosome coordinates. Bars indicate non-coding regions and rectangles indicate coding regions on the transcript models. The signal corresponds to the P site of the codon being translated, and it is shown for all the datasets taken together. **(E)** Schematic showing the coding sequence of the *PHYRE* uORF in the wild-type, *phyre-cr1*, and *phyre-cr2* mutants (top). The start and the stop codons are in bold. The deletion in *phyre-cr1* is indicated by a red dashed line, and the insertion in *phyre-cr2* is highlighted in green. The schematic of the *PHYB* locus is shown in the middle, and the *phyB-cr* alleles are shown at the bottom. The promoter and downstream regulatory regions are shown as black bars, 5’ and 3’ UTRs as grey boxes, and coding region as yellow boxes separated by introns (black dotted lines). The *PHYRE* uORF, *PHYB* start codon, and stop codon are indicated by a magenta rectangle, black arrowhead, and black asterisk, respectively. The binding sites of sgRNAs 1-5, used to generate the *phyre-cr* and *phyB-cr* mutants, are indicated by arrows. Deleted regions in the *phyB-cr* mutants are indicated by red dashed lines. **(F-H)** Representative images and the hypocotyl length quantification of 6-day-old seedlings of the indicated genotypes grown under **(F)** long-day white light or **(G)** long-day deep shade conditions. Scale bars = 5 mm. **(H)** Violin plots show the distribution of individual observations, with overlaid box plots indicating the median and interquartile range. Different letters indicate statistically significant differences among the genotypes (adjusted P < 0.05), determined using the Kruskal-Wallis test followed by Dunn’s post hoc test with Benjamini-Hochberg correction. n = 31-51 and 28-48 seedlings for white light and deep shade treatments respectively.

Among the high-confidence sORFs associated with photomorphogenesis genes, one candidate immediately drew our attention. An uORF residing in the 5′UTR of *PHYB*, the gene encoding the master red-light photoreceptor, was translated under both white light and shade conditions (Figs. 1D, S1C). We named this uORF *PHYTOCHROME REGULATOR* (*PHYRE*). The *PHYRE* stop codon lies only 35 bp upstream of the *PHYB* start codon (Fig. S1D), placing it in unusually close proximity to the downstream ORF and prompting us to investigate whether *PHYRE* translation regulates *PHYB* expression.

To uncover the role of *PHYRE*, we generated a series of frameshift mutants, *phyre-cr,* using a modular cloning-based CRISPR-Cas9 system^27^. These *phyre-cr* mutants either carried a single base insertion or 1-2 base deletion in the *PHYRE* coding sequence, resulting in frameshifts that abolished the native stop codon and led either to premature termination or translational read-through into *PHYB* (Fig. 1E; S2A). In addition, we generated two *phyB-cr* alleles carrying large deletions in the *PHYB* locus that disrupted both *PHYRE* and *PHYB* ORFs (Fig. 1E). We analysed the hypocotyl lengths in white light and deep shade regimes to assess the shade avoidance response of *phyB-cr* and *phyre-cr* mutants. Both *phyB-cr* mutants exhibited elongated hypocotyls under white light and hypocotyl elongation comparable to the wild type under deep shade (Fig. 1F, H), similar to previously reported *phyB* loss-of-function mutants^28,29^. Strikingly, all five independent *phyre-cr* mutants displayed elongated hypocotyls under white light indistinguishable from those of *phyB-cr* mutants (Figs. 1F, 1H; S2B), establishing that *PHYRE* is required to suppress hypocotyl growth under these conditions. In contrast, *phyre-cr* mutants had about 30% shorter-than-wild-type hypocotyls under deep shade (Fig. 1F, H; S2C), indicating *PHYRE* may promote growth specifically in shade. These results suggest that *PHYRE* has opposite roles in regulating hypocotyl growth. In white light it suppresses growth like *PHYB* and in deep shade conditions it promotes growth. Because phyB accumulates in dormant seeds and promotes germination in darkness^30^, we next asked whether *PHYRE* also contributes to this developmental response. We therefore quantified germination efficiency of *phyre-cr* seeds in darkness following a 15-min pulse of red light, which activates phyB, or far-red light, which inactivates it (Fig. 2A). Following red-light treatment, germination efficiency in *phyre-cr* mutants was severely reduced (8–10%), closely resembling *phyB-cr1* (5%) and contrasting sharply with wild-type seeds (75%) (Fig. 2B,E), consistent with the germination defects previously reported for phyB loss-of-function mutants^30,31^.

**Figure 2.**
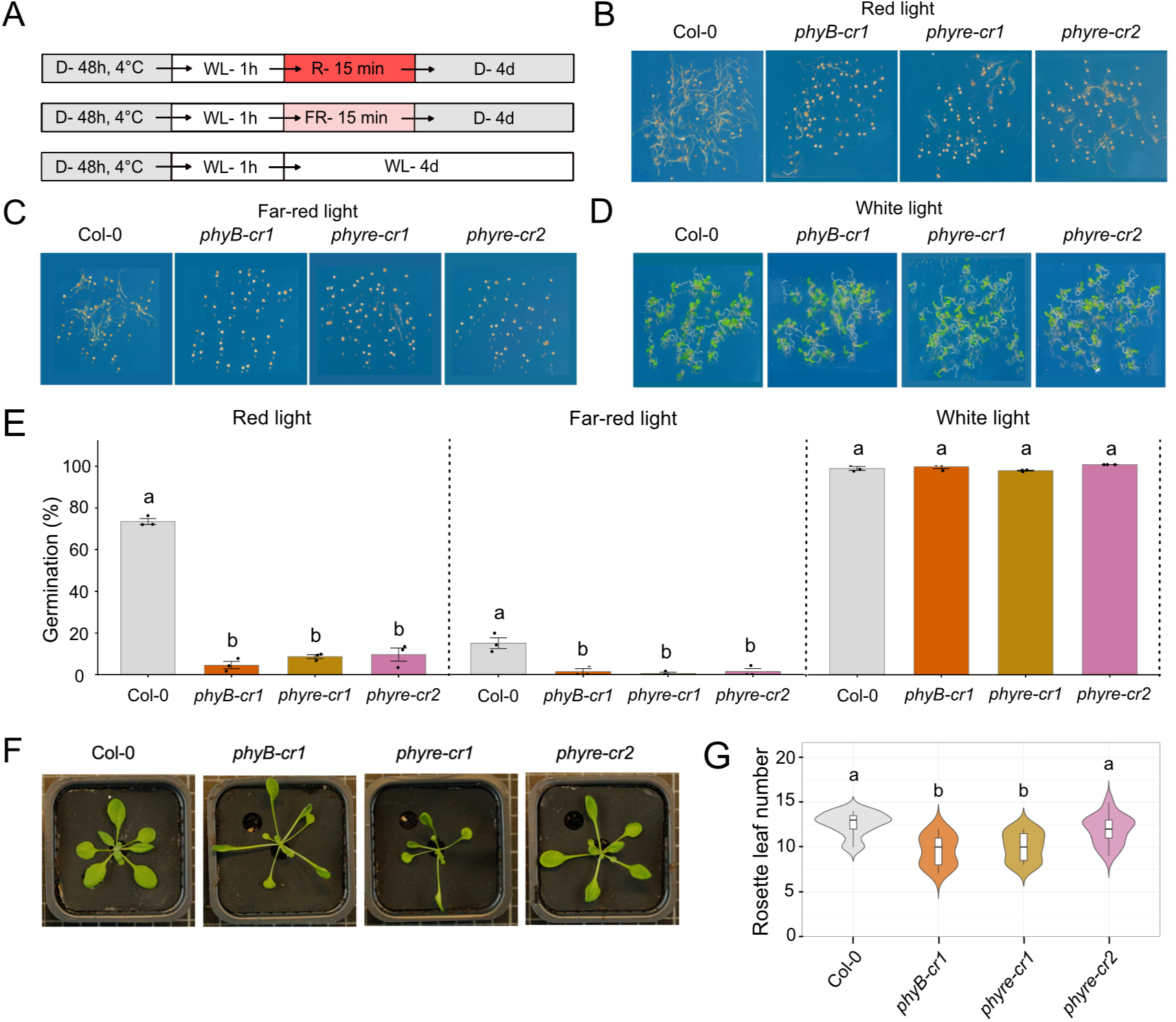
Seed germination, rosette, and flowering time phenotypes of *phyB-cr* and *phyre-cr* mutants. **(A)** Schematic of the light treatment used in the germination experiments. Abbreviations-D = Dark, WL= white light, FR= far-red light, R= red light. **(B-D)** Representative images of seeds/germinated seedlings of the indicated genotypes and light treatments. **(E)** Bar plots showing the mean ± standard error (SE) of the percentage of germination, with individual replicate measurements shown as points, for the indicated genotypes and light treatments. Different letters above the bar plots indicate statistically significant differences among genotypes (adjusted P < 0.05), determined using One-way ANOVA followed by Tukey’s HSD (Honestly Significant Difference) post hoc test. N = 3 (32-75 seeds per biological replicate). **(F)** Representative rosette images of 21 day-old white light-grown plants of indicated genotypes. **(G)** Flowering time quantification indicated by the number of rosette leaves at bolting in the corresponding genotypes on the X-axis. Violin plots show the distribution of individual observations, with overlaid box plots indicating the median and interquartile range. Different letters indicate statistically significant differences among genotypes (adjusted P < 0.05), determined by Kruskal-Wallis test followed by Dunn’s post hoc test with Benjamini-Hochberg correction. n = 13-23 plants.

Following far-red light treatment, germination remained strongly impaired in *phyre-cr* and *phyB-cr* mutants (0.5-1.5%), in contrast to wild-type seeds, which reached 15% germination (Fig. 2C,E). Under white light, germination efficiency exceeded 95% in both *phyre-cr* and *phyB-cr* mutants, indistinguishable from wild type (Fig. 2D,E). Together, these results demonstrate that *PHYRE* is required for efficient seed germination under conditions that depend on phyB activity.

To investigate the impact of *PHYRE* at later developmental stages, we examined rosette development and flowering time in *phyB-cr* and *phyre-cr* mutants grown under long-day white-light conditions. Both *phyre-cr1* and *phyre-cr2* displayed elongated petioles and narrower leaves resembling *phyB-cr1*, although the phenotype of *phyre-cr2* was intermediate between *phyB-cr1* and wild type (Fig. 2F). Consistent with *phyB-cr1*, *phyre-cr1* exhibited early flowering, whereas *phyre-cr2* displayed a more variable flowering-time phenotype that overall remained closer to the wild type (Fig. 2G). Taken together, these findings indicate that *phyre-cr1* and *phyre-cr2* represent strong and weak alleles of *PHYRE*, respectively.

The frameshift mutation in *phyre-cr1* disrupts the normal *PHYRE* reading frame, causing translation to continue into the *PHYB* locus in an alternative frame. Translation terminates after only 31 amino acids because of a premature stop codon, thereby preventing production of any peptide similar to phyB (Fig. S3A). The single-nucleotide insertion in *phyre-cr2* increases the spacing between *PHYRE* and *PHYB* to 36 nt. This eliminates the native *PHYRE* stop codon, thereby enabling complete translational read-through into the *PHYB* ORF and potentially producing an extended phyB protein containing additional extra N-terminal peptide. Given these distinct molecular outcomes, we directly tested whether phyB protein abundance is affected in the two mutants.

We first quantified phyB protein levels in 7-day-old seedlings by western blotting with an anti-phyB antibody. phyB protein was undetectable in *phyre-cr1*, *phyB-9* and *phyB-cr1*, whereas *phyre-cr2* seedlings had reduced levels of phyB under both white-light and shade conditions compared with the wild type (Figs. 3A, S3B). To investigate whether the lower phyB abundance in *phyre-cr* lines could be due to reduced mRNA levels, we determined *PHYB* transcript levels in 7-day old white light-grown seedlings. RT-qPCR analysis with primers targeting N and C-terminus of *PHYB* (Fig. S3C) revealed that the mRNA level was reduced by 50% in the *phyre-cr1* mutant, possibly due to non-sense mediated decay^32^, but remained unchanged in *phyre-cr2* (Fig. 3B). As expected, the *PHYB* transcript was not detected in *phyB-cr1*. The transcript level of *PIL1*, a known gene target repressed by phyB^33^, was increased 10-fold in *phyre-cr1*, similar to *phyB-cr1*, and 2.5-fold in *phyre-cr2*. The reduction in *PHYB* transcript levels in *phyre-cr1* is insufficient to account for the near-complete loss of phyB protein and the strong shade-avoidance phenotype, establishing that *PHYRE* regulates phyB abundance primarily at the translational level.

**Figure 3.**
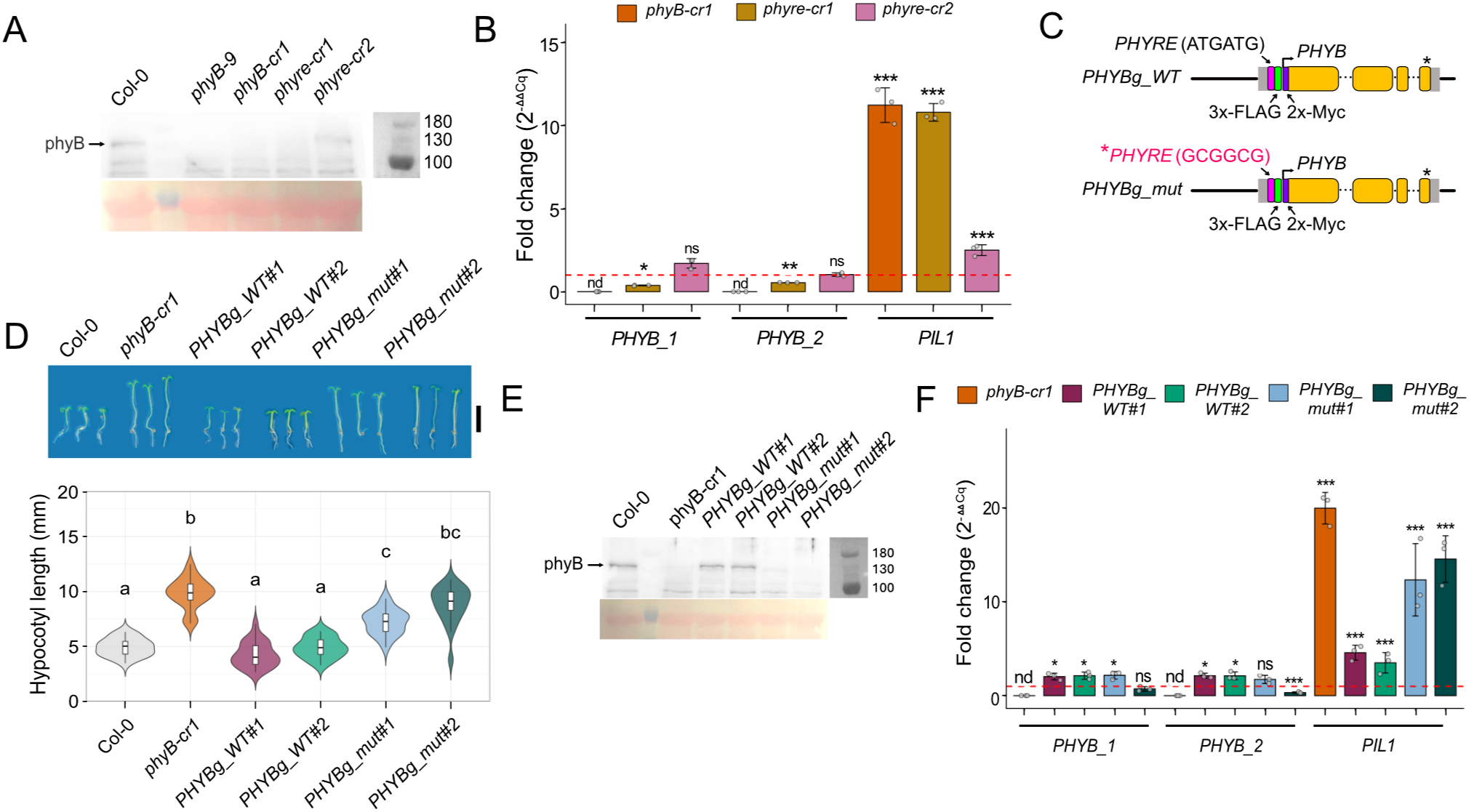
PhyB protein and transcript levels in *phyB-cr*, *phyre-cr* mutants and *PHYB* complementation lines. (A) and. **(E)** Western blot images showing phyB protein levels (129 kDa) in the total protein extracts from 7-day white light-grown seedlings of the indicated genotypes. Image of the corresponding ponceau stained blots (loading control) are shown below. **(B) and (F)** Bar plots showing relative transcript levels of *PHYB* (primer pair-1 and 2) and *PIL1* transcripts in the genotypes indicated in the legend above the plot. Asterisks denote statistical differences of the genotypes compared to wild-type as determined by One-way ANOVA and Dunnett’s post hoc test. * p<0.05; ** p<0.01; *** p<0.001; ns p>0.05, nd. not detected. Error bars denote SD of 3 biological replicates. Red dotted line indicates wild-type transcript levels (normalized to 1). **(C)** Schematics of the wild-type (*PHYBg_WT*) and start codon mutated (*PHYBg_mut*) versions of transgene constructs used for complementing *phyB-cr1* mutant. The promoter and downstream regulatory regions are shown as black bars, 5’ and 3’ UTRs as grey boxes, and coding region as yellow boxes separated by introns (black dotted lines). The *PHYRE* uORF, *PHYB* start codon, and stop codon are indicated by a magenta rectangle, black arrowhead, and black asterisk, respectively. The C-term 3x-FLAG of *PHYRE* and the N-term 2x-Myc tag of *PHYB* are indicated by green and purple rectangles, respectively. **(D)** Representative images and the hypocotyl length quantification of 6-day-old seedlings of the indicated genotypes grown under long-day white light conditions. Scale bar = 5 mm. Violin plots show the distribution of individual observations, with overlaid box plots indicating the median and interquartile range. Different letters indicate statistically significant differences among genotypes (adjusted P < 0.05), determined using Kruskal-Wallis test followed by Dunn’s post hoc test with Benjamini-Hochberg correction. n = 19-23 seedlings.

To test if *PHYRE* translation is required for phyB expression and shade avoidance, we complemented the *phyB-cr1* mutant with *PHYB* genomic fragments expressing either 3x-FLAG-tagged wild-type *PHYRE* or a translation-deficient *PHYRE* variant in which the start codons were mutated, together with 2x-Myc-tagged phyB under the endogenous promoter (Fig. 3C). Two independent lines carrying the wild-type construct (*PHYBg_WT*) fully rescued the hypocotyl elongation phenotype of *phyB-cr1* (Fig. 3D), demonstrating that the transgene was functional. However, two independent lines carrying the translation deficient *PHYRE* genomic fragment (*PHYBg_mut*) failed to restore the hypocotyl length of *phyB-cr1* mutants to wild-type level. Consistent with these phenotypes, phyB protein was restored to wild-type levels in *PHYBg_WT* lines but remained undetectable in *PHYBg_mut* lines (Fig. 3E). As *PHYB* transcript abundance was restored to wild-type levels in both *PHYBg_WT* and *PHYBg_mut* lines (Fig. 3F), the failure of *PHYBg_mut* to express phyB protein cannot be attributed to reduced transcript levels, further demonstrating that *PHYRE* acts at the translational level. *PIL1* expression was lower in *PHYBg_WT* lines (3–4-fold above Col-0, compared with ∼20-fold in *phyB-cr1*) but remained elevated in *PHYBg_mut* lines (12–15-fold above Col-0). The petiole and leaf blade phenotypes of *PHYBg_WT* plants were also similar to those of wild-type plants, whereas *PHYBg_mut* plants largely resembled *phyB-cr1* (Fig. S4A). Similarly, flowering time was restored in *PHYBg_WT* plants, whereas *PHYBg_mut* plants flowered early, as did *phyB-cr1* (Fig. S4B). Together, these results establish that *PHYRE* translation is both necessary and sufficient for phyB protein expression and for normal photomorphogenesis across multiple developmental contexts, demonstrating that translational rather than transcriptional control is the critical regulatory step.

The absolute dependence of phyB protein expression on *PHYRE* translation raised the question of whether this translational control is intrinsic to the *PHYB* 5′UTR or specific to the *PHYB* ORF. To distinguish these possibilities, we replaced the *PHYB* coding sequence with *GUS*, placing it under the control of either a wild-type *PHYRE*-containing 5′UTR (*pPHYB::GUS*) or a start-codon-mutated version (*mutpPHYB::GUS*) driven by the native *PHYB* promoter. GUS activity was undetectable in seedlings carrying *mutpPHYB::GUS* compared with *pPHYB::GUS* controls (Fig. 4A,B), demonstrating that *PHYRE*-dependent translational activation is a general property of the *PHYB* 5′UTR and not specific to the *PHYB* coding sequence.

**Figure 4.**
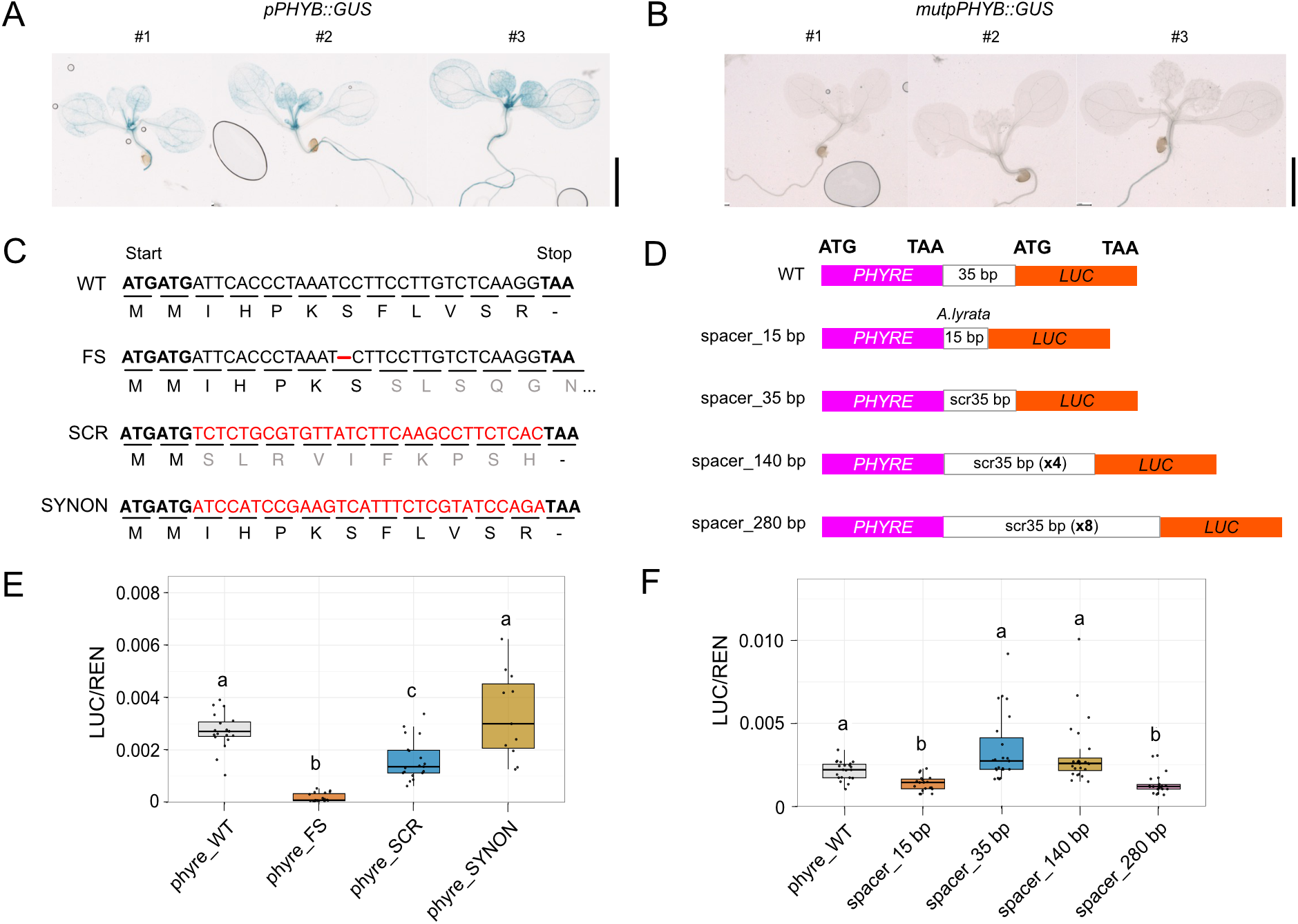
Results of GUS and luciferase reporter assays for *PHYB* 5’ UTR variants. **(A-B)** Representative images of GUS-stained 7-day-old seedlings from three independent lines each of **(A)** *pPHYB::GUS* (wild type) and **(B)** *mutpPHYB::GUS* (*PHYRE* start codon mutant) genotypes. Scale bar = 2 mm. **(C)** Schematic representation of the DNA and the amino acid sequence of the *PHYRE* variants used for lucifrase reporter assays in **(E)**. WT-wild type, FS-frameshift, SCR-scrambled, SYNON-synonymous mutation. Deletion in FS is indicated by red dashed line. Changes in coding sequence in SCR and SYNON constructs are indicated in red. Changes in amino acid sequence in FS and SCR constructs are indicated in grey. **(D)** Schematic of the spacer variants used for luciferase reporter assays in **(F)**. Magenta and orange rectangles represent coding sequences of *PHYRE* and firefly luciferase (*LUC*) respectively. **(E-F)** Quantification of relative firefly luciferase activity in tobacco leaf extracts infiltrated with the indicated reporter constructs. Box plots indicate the median and interquartile range (IQR), with whiskers extending to 1.5 × IQR. Individual points represent leaf discs. Different letters above the violin plots indicate statistical differences among reporter constructs (adjusted P < 0.05), determined using Kruskal-Wallis test followed by Dunn’s post hoc test with Benjamini-Hochberg correction. n= 12-24 leaf discs from 4-8 mature leaves. WT-wild type, FS-frameshift, SCR-scrambled, SYNON-synonymous mutation.

uORFs regulate downstream translation through three distinct mechanisms: nucleotide sequence, nascent peptide identity, and intercistronic distance ^34,35,36^. To distinguish whether the regulatory activity of *PHYRE* resides in its RNA sequence or encoded peptide, we measured firefly luciferase activity from reporter constructs carrying *PHYB 5′UTRs* with frameshift, scrambled, or synonymous mutations in *PHYRE* (Fig. 4C). Scrambling the *PHYRE* sequence altered the encoded amino acid sequence whereas synonymous mutations altered the nucleotide sequence without changing the encoded peptide sequence. The constructs were driven by the native Arabidopsis *PHYB* promoter and transiently expressed in tobacco leaves. Relative to the wild-type *PHYRE* control, a frameshift identical to that in *phyre-cr1* caused a marked reduction in luciferase activity, while scrambling the PHYRE peptide sequence resulted in a partial reduction (Fig. 4E). Synonymous recoding, which preserved the wild-type peptide while altering the RNA sequence, had no effect on luciferase activity. Together, these results indicate that the PHYRE peptide sequence, rather than the RNA sequence, contributes to downstream translational activation.

To disentangle the effects of spacer sequence and length on downstream ORF expression, we scrambled the native 35-bp region separating the *PHYRE* stop codon from the downstream *PHYB* start codon and generated longer spacers of 140 and 280 bp by tandemly repeating the scrambled sequence, ensuring that no additional ORFs were introduced (Fig. 4D). We also generated a construct with a 15 bp spacer matching the shorter intercistronic distance found in *Arabidopsis lyrata*, whose *PHYRE* sequence is otherwise identical to that of *Arabidopsis thaliana*. The reporter carrying the scrambled 35 bp spacer exhibited luciferase activity comparable to that of the wild-type control (Fig. 4F), indicating that spacer sequence has little influence on downstream ORF expression. In contrast, shortening the spacer to 15 bp or extending it to 280 bp significantly reduced luciferase activity. Interestingly, extending the spacer to 140 bp did not affect luciferase activity, which remained comparable to that of the 35-bp spacer control. Together, these findings establish that intercistronic distance is a key determinant of *PHYRE*-dependent translational activation, pointing to a position-sensitive mechanism linking uORF translation to downstream ORF expression.

## DISCUSSION

*PHYRE* defines one of the few characterised activating uORFs, and our results establish its translation as an essential prerequisite for phyB protein expression and normal photomorphogenesis across multiple developmental contexts (Fig. 1-3). Disrupting *PHYRE* translation through either a frameshift or start codon mutation eliminates phyB protein expression without substantially affecting *PHYB* mRNA levels (Fig. 3), suggesting that *PHYRE* acts primarily at the translational level. We propose that translation of *PHYRE* is required for translation of the downstream *PHYB* ORF (Fig. 5). The spacer experiments indicate that efficient *PHYB* translation depends on the distance between the *PHYRE* stop codon and the *PHYB* start codon, with both shorter and longer spacers reducing downstream translation (Fig. 4). These findings support a model in which translation of *PHYRE* facilitates reinitiation at the downstream *PHYB* start codon, with the efficiency of this process depending on the intervening distance.

**Figure 5:**
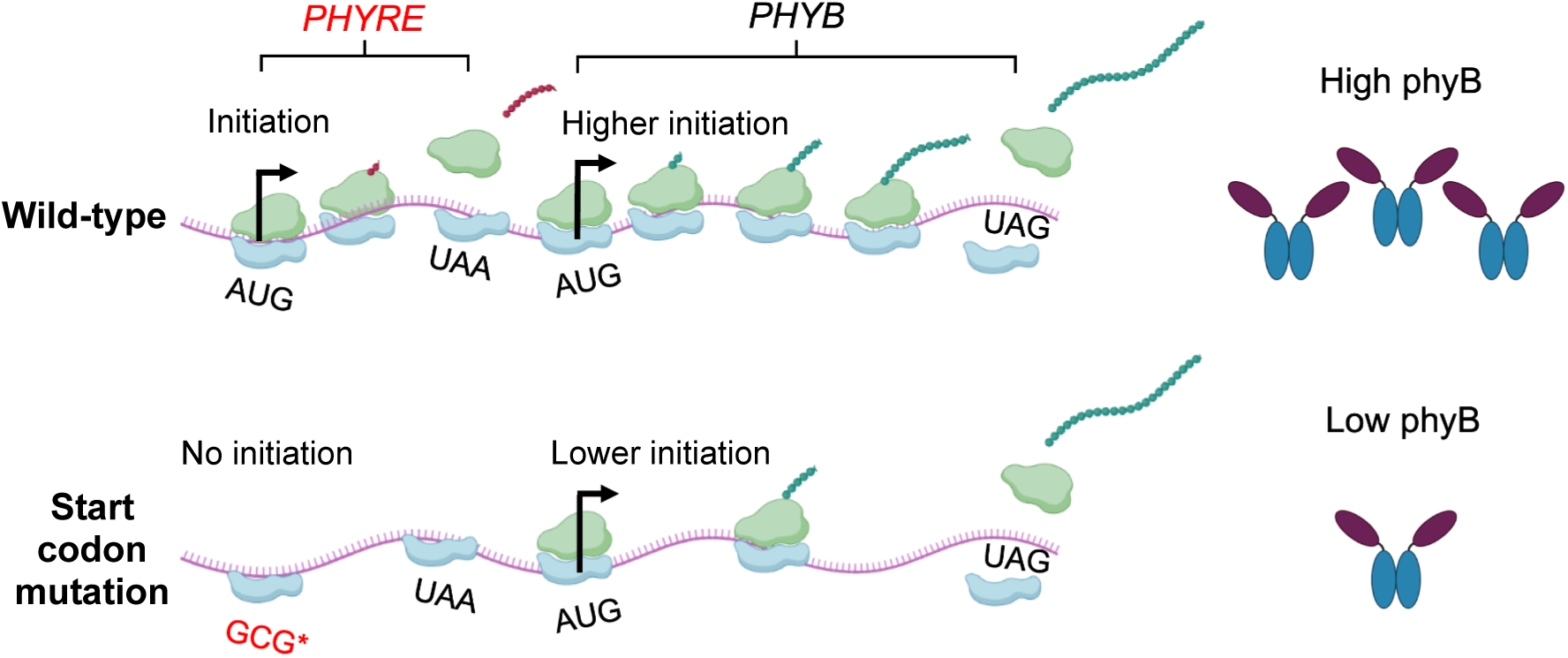
Proposed model for *PHYRE*-mediated regulation of *PHYB* translation. Position-dependent *PHYRE* uORF translation maintains optimal phyB protein expression, possibly by facilitating translation re-initiation at the *PHYB* start codon. Impaired *PHYRE* translation, for example due to mutation of its start codon, results in reduced phyB protein levels, possibly because ribosomes bypass *PHYRE* and continue scanning, thereby reducing translation initiation at the downstream *PHYB* start codon.

Whether position-dependent uORF-mediated regulation of *PHYB* translation is evolutionarily conserved is an important question. A survey of the 150 bp genomic region upstream of the *PHYB* coding sequence across 49 land plant species revealed short proximal uORFs of 5–20 aa across eudicots, while monocot and gymnosperm representatives lacked such elements entirely (Fig. S5). In *Brassicaceae*, a proximal uORF is maintained at a similar relative position across major genera from *Eutrema* to *Brassica*, consistent with functional constraint on uORF position. However, peptide-level sequence conservation is restricted to species closely related to *Arabidopsis thaliana*, suggesting that across eudicots more broadly, the act of uORF translation rather than the identity of the encoded peptide may be the primary conserved regulatory feature. Whether divergent PHYRE peptides share a conserved three-dimensional structure that underlies their function across species is an intriguing question that warrants further investigation. Together, these observations are consistent with a model in which uORF-mediated translational activation of *PHYB* is a eudicot innovation, with peptide-level contributions to this regulation arising more recently within the *Brassicaceae*.

The partial reduction in luciferase activity observed with a scrambled PHYRE peptide sequence (Fig. 4) supports the idea that peptide identity contributes to the efficiency of downstream translational activation in *Arabidopsis thaliana*, consistent with the more restricted conservation of the PHYRE peptide sequence within the *Brassicaceae*. Previous studies have shown that uORF-encoded peptides can regulate downstream ORF translation in genes involved in metabolic processes, including *bZIP11* and *Polyamine Oxidase2*^37,38^. A well-characterised precedent is the sucrose control peptide encoded by a uORF in the *bZIP11* 5′UTR, which induces ribosome stalling under high-sucrose conditions to inhibit downstream translation. *PHYRE* represents a mechanistic inversion of this paradigm, promoting rather than inhibiting downstream translation, yet whether this activity is mediated through direct peptide-ribosome interactions during translation or termination, or simply through the act of uORF translation itself, remains to be established.

An intriguing observation is that *phyre-cr* mutants consistently exhibited shorter hypocotyls than wild-type plants under deep shade, a condition under which phyB activity is strongly reduced and *phyre-cr* mutants are therefore expected to phenocopy *phyB-cr* lines (Fig. 1). This difference raises the possibility that the PHYRE peptide has an additional role in regulating hypocotyl elongation under shade, independent of its function in promoting phyB translation.

Why would plants evolve *PHYRE*-mediated translational control of a constitutively transcribed photoreceptor? phyB, along with phyA, contribute to shade perception and the regulation of growth under changing light conditions, with phyB playing a prominent role in sensing changes in the R:FR ratio^39,40^. Moreover, phyB acts as both a photoreceptor and a thermosensor, integrating light and temperature signals to regulate plant growth and development^41,42^. Recent evidence indicates that phyB abundance influences photobody formation and Phytochrome Interacting Factor (PIF) activity, thereby shaping downstream signalling^43^. Thus, *PHYRE*-mediated translational control provides an additional mechanism for tuning phyB abundance as environmental conditions change. The detection of *PHYRE* translation under both white light and shade indicates that this mechanism operates continuously across distinct light regimes, positioning it to act as a constitutive translational amplifier of phyB output. Such regulation is likely to be particularly important when rapid shifts in light quality or temperature alter the balance between active and inactive phyB, requiring proportionate adjustments in photoreceptor abundance that cannot be achieved through transcriptional regulation of a constitutively expressed gene.

## MATERIALS AND METHODS

### Plant materials and growth conditions

*Arabidopsis thaliana* ecotype Columbia-0 (Col-0) was used as the wild-type control. All the lines used in this study including *phyB-9;ven4+/+* (Reed et al.^44^; Gift from Dr. Andreas Hiltbrunner, University of Freiburg) are in Col-0 genetic background. Seeds were surface sterilized using sodium hypochlorite (1.4%) in 70% ethanol containing 0.001% TritonX-100 for 2-3 minutes. Surface sterilized seeds were spotted on plates with 0.5x MS medium including vitamins (Duchefa Biochemie M0222) supplemented with 0.1% Sucrose (Duchefa Biochemie S0809), and 0.8% agar (Duchefa Biochemie P1001). Seeds were stratified for 3 days in dark at 4°C and shifted to Percival LED growth chamber (Model LED-30SB10) for seed germination, hypocotyl growth, and ribosome profiling experiments.

### DNA constructs

Genome editing constructs for *phyB-cr* and *phyre-cr* generation were assembled using the modular cloning (MoClo) approach described in Grutzner et al.^45^; Stuttmann et al.^27^ Briefly, sgRNAs targeting *PHYB* or *PHYRE* were designed using CRISPR-P 2.0 (http://crispr.hzau.edu.cn/CRISPR2/) and cloned individually in shuttle vectors to generate sgRNA TUs. Multiple sgRNA TUs were assembled in pDGE347 recipient binary vector containing zCas9i and FAST selection cassettes (zCas9i cloning kit, Addgene Kit # 1000000171). Protocols for MoClo assembly reactions were followed exactly as described in zCas9i cloning manual v5 from Addgene Kit # 1000000171.

*PHYBg_WT* and *PHYBg_mut* complementation constructs were assembled using NEBuilder ® HiFi DNA Assembly Reaction Protocol (NEB #E2621) according to manufacturer’s instructions. Briefly, synthetic fragments containing C-term 3x-FLAG tagged wild-type or start-codons mutated version of *PHYRE* and N-term 2x-Myc tagged *PHYB* region (Twist Bioscience) was assembled with gel purified PCR fragments spanning 7.2 kb *PHYB* intergenic region into pFAST-R07 (https://vectorvault.vib.be/collection/pfast-r07) binary vector containing FAST cassette for seed coat RFP selection.

For generating *pPHYB::GUS* and *mutPHYB::GUS* constructs, 2 kb *PHYB* promoter was assembled along with 300 bp 5’UTR containing wild-type or start-codons mutated version of *PHYRE* and *GUS* coding sequence into the gateway multisite destination vector pH7m34GW (https://vectorvault.vib.be/collection/ph7m34gw) using Gateway LR Clonase II enzyme mix (Invitrogen).

Dual luciferase reporter constructs were generated by GreenGate cloning system. *PHYB* 5’UTR variants were synthesized from Twist bioscience and assembled along with 2 kb *PHYB* promoter, firefly luciferase coding sequence and p35S renilla luciferase modules into the binary destination vector pGGZ003 (GreenGate Cloning System, Addgene Kit # 1000000036). Protocols for GreenGate assembly reactions were set up as described in Lampropoulos et al.^46^ Gene fragments were assembled in single cut/ligation reactions containing 1μl of FastDigest Eco31I (Thermo Scientific FD0293), 15 U of T4 DNA Ligase, HC (Thermo Scientific EL0013) and 1mM ATP (Thermo Scientific R0441).

Genome editing, complementation, and GUS constructs were transformed into *Agrobacterium tumefaciens* GV3101, while dual luciferase constructs were transformed into *Agrobacterium tumefaciens* GV3101 (pSoup) strain. Colonies were screened for insert by PCR before Col-0 transformation. Oligos used for cloning are included in the Supplementary table 3.

### Plant transformation and line establishment

*Agrobacterium tumefaciens* carrying DNA constructs were transformed into 4 week old *Arabidopsis thaliana* Col-0 plants by floral dipping as described in Clough and Bent^47^.The transformants were screened for seed coat RFP fluorescence under Leica MZ FLIII fluorescence stereomicroscope fitted with CoolLED pE-300^lite^ illumination system. For *phyB-cr* and *phyre-cr* line establishment, RFP negatives were selected from T2 segregating population and genotyped for deletions and point mutations by PCR, followed by sanger sequencing. Positive T2 plants were selfed to establish T3 homozygous lines. T3 lines were used for all the experiments described in this study. For *PHYBg_WT* and *PHYBg_mut* complementation, T1 RFP positive plants were selected and transgene insertion was confirmed by PCR. T2 segregating population obtained from T1 RFP positive plants was used for qPCR and western blotting experiments. For hypocotyl assays under white light and deep shade, RFP positives (homozygous/heterozygous for transgene) were selected from T2 segregating population and spotted on 0.5X MS plates. For GUS assays, hygromycin resistant T1 plants were selected on 0.5X MS agar plates supplemented with 25μg/ml hygromycin B (Duchefa Biochemie H0192) and 100 μg/ml cefotaxime sodium (Duchefa Biochemie C0111). Insertion of GUS-transgene in hygromycin resistant T1 plants was verified by genotyping PCR. T2 segregating population obtained from T1 positive plants was used for GUS assay. All genotyping PCR primers are listed in Supplementary table 3.

### Hypocotyl measurements

Following 3 days of stratification in dark at 4°C, plates were shifted to Percival LED growth chamber (Model LED-30SB10) set under white light condition (WL; PAR-6.5 μmolm^-2^s^-1^; R:FR= 12.1; 16-h light/8-h dark photoperiod) at 22°C, 65% relative humidity for 2 days for germination and establishment. Seedlings were grown further for 4 days under white light or deep shade (WL+ FR; PAR-6.5 μmolm^-2^s^-1^; R:FR= 0.15; 16-h light/8-h dark photoperiod) conditions at 22°C and 65% relative humidity. 6-day old seedlings were imaged using Epson scanner expression 12000XL equipped with SilverFast9 (Laser Software Imaging). Hypocotyls were measured using segmented line tool from ImageJ v1.54t software. Representatives images were compiled by removing background using online background remover tool from Canva and performing brightness/contrast adjustments using in-built Microsoft Powerpoint function.

### Seed germination

Seed germination assays were performed as described in Gu et al.^31^ with minor modifications. Following 3 days of stratification in dark at 4°C, plates were placed in Percival LED growth chamber (Model LED-30SB10) under white light (PAR-100 μmolm^-2^s^-1^; R:FR= 12.1) for 1 hour at 22°C, 65% relative humidity, followed by 15 min of monochromatic red light (35 μmolm^-2^s^-1^) or far-red light (38 μmolm^-2^s^-1^) treatment. After light treatment, plates were maintained at 22°C in darkness for 4 days before germination rates were calculated.

### Flowering time

6 day-old white light grown seedlings (PAR-6.5 μmolm^-2^s^-1^; R:FR= 12.1; 16-h light/8-h dark photoperiod) were transferred to pots containing a 3:1 soil:vermiculite mixture (Hasselfors garden specialjord; Sibelco vermiculite) supplemented with 1% SW Bouyant RIKA S 7-1-5 fertilizer and nematode mixture (NEMAblom; BioNEMA). Pots were maintained in invitro growth rooms under white light irradiation (PAR-180 μmolm^-2^s^-1^; 16-h light/8-h dark photoperiod) at 22°C, 65% relative humidity until bolting. Flowering time was determined by counting the number of rosette leaves when the bolt reached a height of 5 cm.

### Ribosome profiling and library preparation

Seven-day-old white light (WL; PAR-6.5 μmolm^-2^s^-1^; R:FR= 12.1; 16-h light/8-h dark photoperiod) grown seedlings were subjected to deep-shade treatment (WL+ FR; PAR-6.5 μmolm^-2^s^-1^; R:FR= 0.15) for 45 or 90 min, while control seedlings were maintained under white light for an equivalent 90-min period. All samples were harvested simultaneously at the end of the 90-min experimental period and immediately flash-frozen in liquid nitrogen. Frozen tissue was ground to a fine powder in liquid nitrogen, and ribosome-protected fragments (RPFs) were isolated and processed for ribosome profiling (Ribo-seq) as described in Mahboubi et al^48^. Briefly, powdered tissue was extracted in ribosome extraction buffer containing cycloheximide, and the lysates were digested with RNase I to remove unprotected RNA. RPFs were purified by TRIzol extraction and isopropanol precipitation, size-selected (27–31 nt) by denaturing polyacrylamide gel electrophoresis, and used for library preparation.

Libraries were constructed by 3′ linker ligation, reverse transcription, cDNA circularization, rRNA depletion, and PCR amplification as described in Mahboubi et al^48^. Purified libraries were quantified and subjected to single-end sequencing.

### Riboseq analysis

Raw Ribo-seq libraries were processed independently. Adapter trimming and quality control were performed separately for each library, followed by alignment to the *Arabidopsis thaliana* reference genome using STAR and the Araport11 annotation^49^. P-site offsets were estimated independently for each library and read length using the bioconductor package ribosomeProfilingQC 1.23.0 (https://bioconductor.org/packages/release/bioc/html/ribosomeProfilingQC.html).

After offset correction, the alignments were pooled into a single coordinate-sorted BAM file containing 126,198,671 primary mapped P-site alignments. Candidate ORFs were evaluated using RibORF v.2.0^26^. We employed a sensitive detection strategy (see Fig. S1) and considered ORFs with a RibORF score of at least 0.6 to show evidence of translation. The final non-canonical ORF catalogue was restricted to uORFs, dORFs and ORFs located in transcripts annotated as non-coding (ncRNA ORFs). Candidates overlapping annotated coding sequences on the same strand were excluded. Redundant predictions sharing the same chromosome, strand and genomic ORF boundaries were collapsed. We identified 2,856 novel translated sORFs using this procedure.

We applied a subsequent filter to focus on a set of 208 highly translated sORFs. These sORFs had to be supported by more than 30 reads and show high signal homogeneity along the ORF (more than 30% of codons covered by reads) and display >50% periodicity.

### Western blotting

Total protein was extracted from finely ground tissue of 7-day-old seedlings grown under white light or deep shade using 100 µL of extraction buffer (50 mM Tris-HCl pH 7.5; 150mM NaCl; 1mM EDTA pH 8.0; 10%, glycerol; 2mM PMSF; 1% Triton X-100; 5 mM DTT; 1x cOmplete™ Protease Inhibitor Cocktail). Prior to separation on SDS PAGE, extracts were quantified by Bradford assay and 60 µg of total protein was resuspended in 1X loading dye (0.05% bromophenol blue, 0.1 M DTT, 2% glycerol, 2% SDS, 0.05M Tris-HCl pH 6.8) and incubated for 5 min at 95°C. Samples were separated on 9% SDS-PAGE and blotted to PVDF membrane (Thermoscientific 88518; pore size 0.45 µm). Blot was blocked with 5% milk powder in TBS-T for 1 h/RT with agitation. Blot was incubated in the primary antibody (Anti-PhyB, Agrisera AS21 4566) at a dilution of 1:1000 in TBS-T ON/4°C with agitation. The antibody solution was decanted and the blot was rinsed with TBS-T briefly twice, then washed 3 times for 10 min in TBS-T at RT with agitation. Blot was incubated in matching secondary antibody (anti-rabbit IgG horse radish peroxidase conjugated, Agrisera AS09 602) diluted to 1:25 000 in TBS-T for 1h/RT with agitation. The blot was washed as above, developed for 5 min with ECL™ Western Blotting Reagents (Cytiva RPN2106) and imaged using C600 Azure biosystems cSeries luminescence camera system under auto exposure settings. Representative blot images were compiled after performing brightness/contrast adjustments using in-built Microsoft Powerpoint function.

### RT-qPCR

Total RNA was extracted from 100mg finely ground tissue of 7-days old white light grown seedlings using RNAeasy Mini extraction kit (Qiagen 74104) according to manufacturer’s protocol. 1µg of DNAse free RNA was converted to cDNA using iScript cDNA synthesis kit (Biorad 1708891) according to manufacturer’s instructions. qPCR was performed in Biorad CFX384 Touch Real-Time PCR Detection System using 1.2µL of cDNA reaction template with 0.5µM primers in 1x KAPA SYBR FAST qPCR Master Mix (KK4607). Fold change (2^-ΔΔCq^) in transcript levels was determined using in-built analysis function of the Click-qPCR tool (Kubota and Tajima.^50^). qPCR primers are listed in Supplementary table 3.

### GUS assay

Samples were incubated in GUS staining solution (50 mM sodium phosphate buffer pH 7.2, 1 mM K_3_Fe(CN)_6_, 1 mM K_4_Fe(CN)_6_, 0.1% Triton X-100, 1 mM X-gluc) for 4h at 37°C with gentle shaking in the dark. Following staining, samples were fixed in FAA (50% ethanol, 5% glacial acetic acid, 5% formaldehyde) for 10 min at room temperature and decolorized in 100% ethanol with gentle shaking, replacing the ethanol as needed until chlorophyll was removed. Samples were then rehydrated in 100%, 90%, and 80% ethanol for 5 min each, incubated in clearing solution I (20% methanol and 0.24 M HCl) for 15 min at 57°C, and transferred to clearing solution II (7% NaOH in 60% ethanol) for 15 min at room temperature. Samples were subsequently incubated in 50% ethanol overnight and stored in 70% ethanol until imaging. Before imaging, samples were rehydrated in 40%, 20%, and 10% ethanol for 5 min each. GUS-stained seedlings were imaged using Leica M205 FCA fluorescence stereo microscope equipped with Leica DFC9000 GT camera and LAS X image analysis software. Images were processed by removing background and adjusting brightness/contrast using in-built Microsoft Powerpoint function.

### Dual Luciferase assay

The reporter constructs were transformed into *Agrobacterium tumefaciens* strain GV3101(pSoup). A single positive colony was cultured overnight in 5 mL LB medium containing the appropriate antibiotics at 28°C. The culture was diluted 1:100 into 50 mL fresh LB medium supplemented with antibiotics and grown overnight at 28°C. Cells were resuspended in infiltration buffer (10 mM MgCl₂, 10 mM MES, pH 5.6, and 150 µM acetosyringone) to an OD₆₀₀ of 0.8 and incubated in the dark for 2 h. Fully expanded leaves of 4-5-week-old *Nicotiana benthamiana* plants were infiltrated using a needleless syringe. For each construct, two leaves from each of two plants (four leaves total) were infiltrated. Plants were maintained at 22°C under a 16-h light/8-h dark photoperiod, with low light and covering for the first 24 h after infiltration. After 72 h, twelve leaf discs (5 mm diameter) were collected from the four infiltrated leaves per construct and stored individually in liquid nitrogen. Non-infiltrated leaf discs were collected as controls for background luminescence.

Each leaf disc was ground individually in liquid nitrogen, and 100 µL of 1x Passive Lysis Buffer (Promega) was added. Lysates were incubated at room temperature for 15 min and centrifuged at 16,000 x g for 20 min at 4°C. Firefly and Renilla luciferase activities were measured using the Dual-Luciferase Reporter Assay System (Promega, E1960) according to the manufacturer’s instructions. For each sample, 10 µL of lysate was assayed in three technical replicates using a GloMax Navigator luminometer (Promega, Madison, WI, USA) with default settings. Background luminescence from non-infiltrated leaf discs was subtracted, and relative promoter activity was calculated as the ratio of firefly (LUC) to Renilla (REN) luciferase activity. The experiment was repeated twice.

### PHYB 5’UTR survey

Genomic sequences 150 nucleotides immediately upstream of the *PHYB* translation start site were retrieved for 49 land plant species spanning gymnosperms, monocots, basal eudicots, asterids, and rosids, with emphasis on *Brassicaceae*. For each species, a RefSeq mRNA accession for *PHYB* was used as the entry point: the corresponding NCBI Gene record was retrieved via the Entrez API (Biopython 1.8x), and the chromosomal accession and gene coordinates were obtained from the Gene esummary. The annotated gene locus plus 350 bp of upstream padding was then fetched from the chromosomal record, the *PHYB* coding sequence (CDS) feature was identified by keyword matching of gene/product annotations (keywords: phyb, phytochrome b, phytochrome-b and variants), and exactly 150 bp immediately upstream of the CDS ATG was extracted, with strand orientation handled by reverse-complementing minus-strand loci. For species where the genomic CDS feature could not be resolved automatically, the analysis fell back to the annotated 5′ UTR of the mRNA record. The 150 bp window was scanned in-frame for all ATG codons followed by in-frame stop codons (TAA, TAG, TGA) within the window; uORF positions are reported as nucleotide offsets relative to the main PHYB ATG. A uORF was classified as *PHYRE*-like if it encoded 5-20 amino acids and its stop codon terminated ≤ 100 nt upstream of the *PHYB* ATG, matching the criteria of the *Arabidopsis thaliana PHYRE* element (12 aa, gap of 38 nt). Tandem ATG pairs (two ATGs separated by ≤ 6 nt) were also recorded. The Arabidopsis *PHYRE* uORF position and sequence (MMIHPKSFLVSR, ATG at −77, stop end at −39) were taken from TAIR. All scripts were written in Python 3 using Biopython V.1.88.

### Data visualization and statistical analysis

For hypocotyl, germination, flowering time and luciferase assays, plots and statistical analysis were conducted using custom R-scripts (R version 4.4.1) in RStudio (Version 2026.09.0+174). For qPCR data, Click-qPCR tool (Kubota and Tajima.^50^) was used to generate bar plots and perform statistical analysis. Details about the statistical analysis are as follows:

For hypocotyl measurements, flowering time, and luciferase assays, statistical differences were assessed using one-way ANOVA followed by Tukey’s HSD test when all groups satisfied the Shapiro-Wilk normality criterion (P > 0.05); otherwise, the Kruskal-Wallis test followed by Dunn’s test with Benjamini-Hochberg adjustment was used. For germination data, statistical differences were assessed by one-way ANOVA followed by Tukey’s HSD test. For qPCR data, first, a one-way ANOVA was performed on the ΔCq values to determine if there are any significant differences among the means of all selected genotypes. Then, Dunnett’s test was used as a post-hoc test to specifically compare each genotype against the wild-type control. All raw data are included in Supplementary table 4.

## Declaration of generative AI and AI-assisted technologies in the manuscript preparation process

During the preparation of this work, we used ChatGTP, Claude and copilot for language editing. The author(s) reviewed and edited the output as needed and take full responsibility for the content of the published article.

## ACKNOWLEDGEMENTS

Financial support was provided by the European Research Council (ERC) Synergy grant RESYDE (101167406), the NovoCrops Centre (Novo Nordisk Foundation grant 2019OC53580), the Novo Nordisk Foundation (grants NNF18OC0034226, NNF20OC0061440, and NNF23OC0084195), the Independent Research Fund Denmark (grants 0136-00015B and 0135-00014B), and Vetenskapsrådet (VR; grant 2024-04792), all to S.W. Additional support was received from the Wallenberg Initiatives in Forest Research (WIFORCE), funded by the Knut and Alice Wallenberg Foundation. We are grateful to Jaume Martínez-García for his insightful comments and critical reading of the manuscript.

## AUTHOR CONTRIBUTIONS

N.S. drove the experimental work and wrote the manuscript. A.V.H., M.J., A.E. and A.Ma. performed experiments. A.Ma. and J.H. contributed to data analysis. J.C.M., C.P. and M.M.A. performed bioinformatic data analysis. S.W. contributed to data analysis, wrote the manuscript, supervised the project, and acquired funding. All authors commented on the manuscript.

---

**Figure S1.**
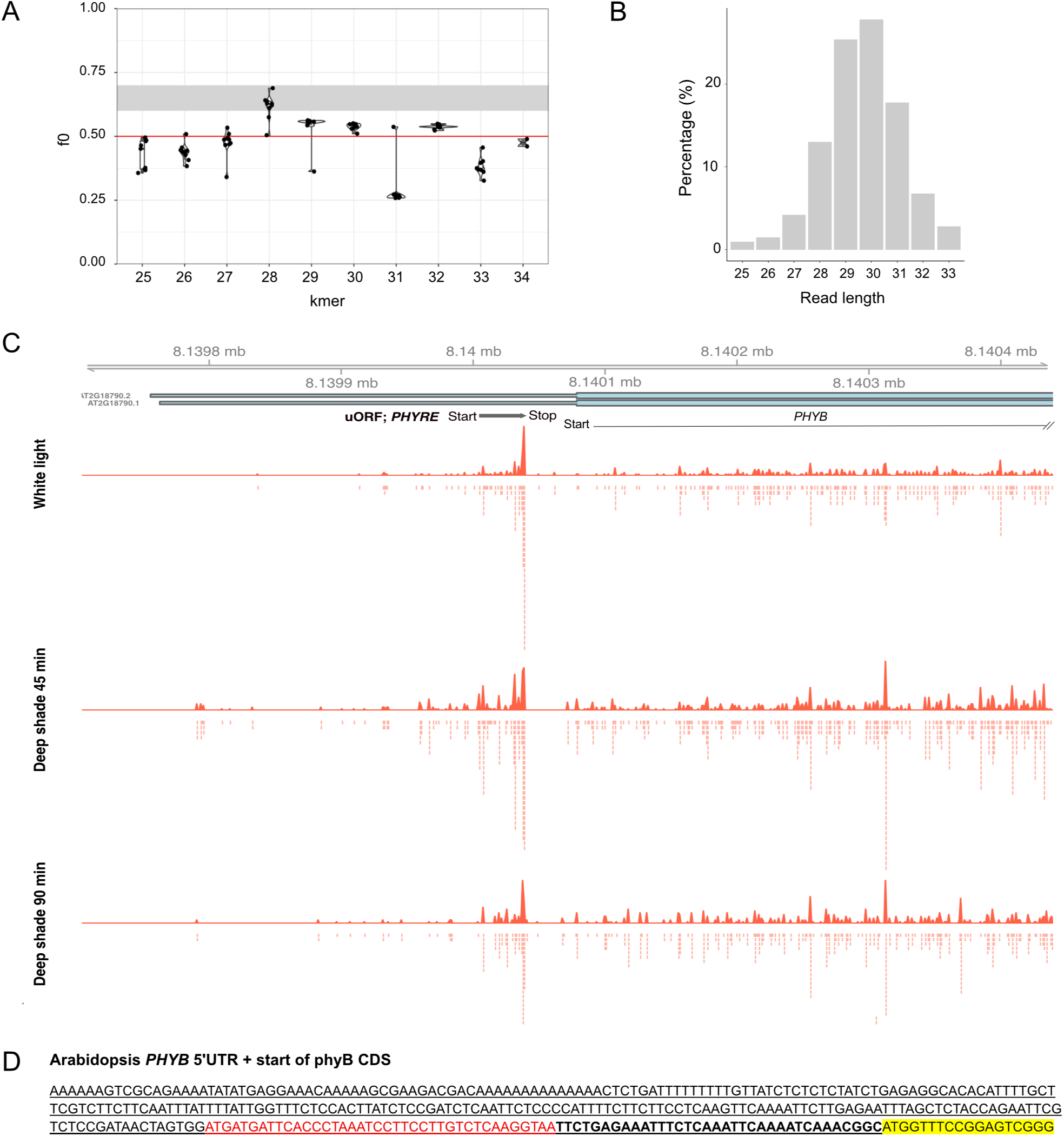
Results from the riboseq analysis. **(A)** Periodicity of reads of different length for the different experiments. The fraction of reads of a given read length (kmer) in the correct frame, in the different experiments, is indicated as f0. A f0 value higher than 0.5, indicating that the majority of reads are in the correct frame, was used as a cut-off to select a given read length in an experiment. **(B)** Read length distribution. The plot shows the percentage of reads of different length, considering only reads that passed the threshold indicated in **(A)**. **(C)** Ribosome profile of *PHYRE* and start of the *PHYB* coding region under different conditions. Transcript gene models in blue are shown below the chromosome coordinates. Bars indicate non-coding regions and rectangles indicate coding regions on the transcript models. The signal corresponds to the P site in the codon that is being translated. The accumulated signal from the three replicates of each condition (white light, deep shade 45’ and deep shade 90’) is shown. **(D)** Sequence information of 334 bp region of *PHYB* gene including the annotated 5’UTR (underlined) and the start of the phyB coding sequence (highlighted in yellow). *PHYRE* coding sequence is in red and the 35 base spacer separating the *PHYRE* stop codon and the *PHYB* start codon is in bold.

**Figure S2.**
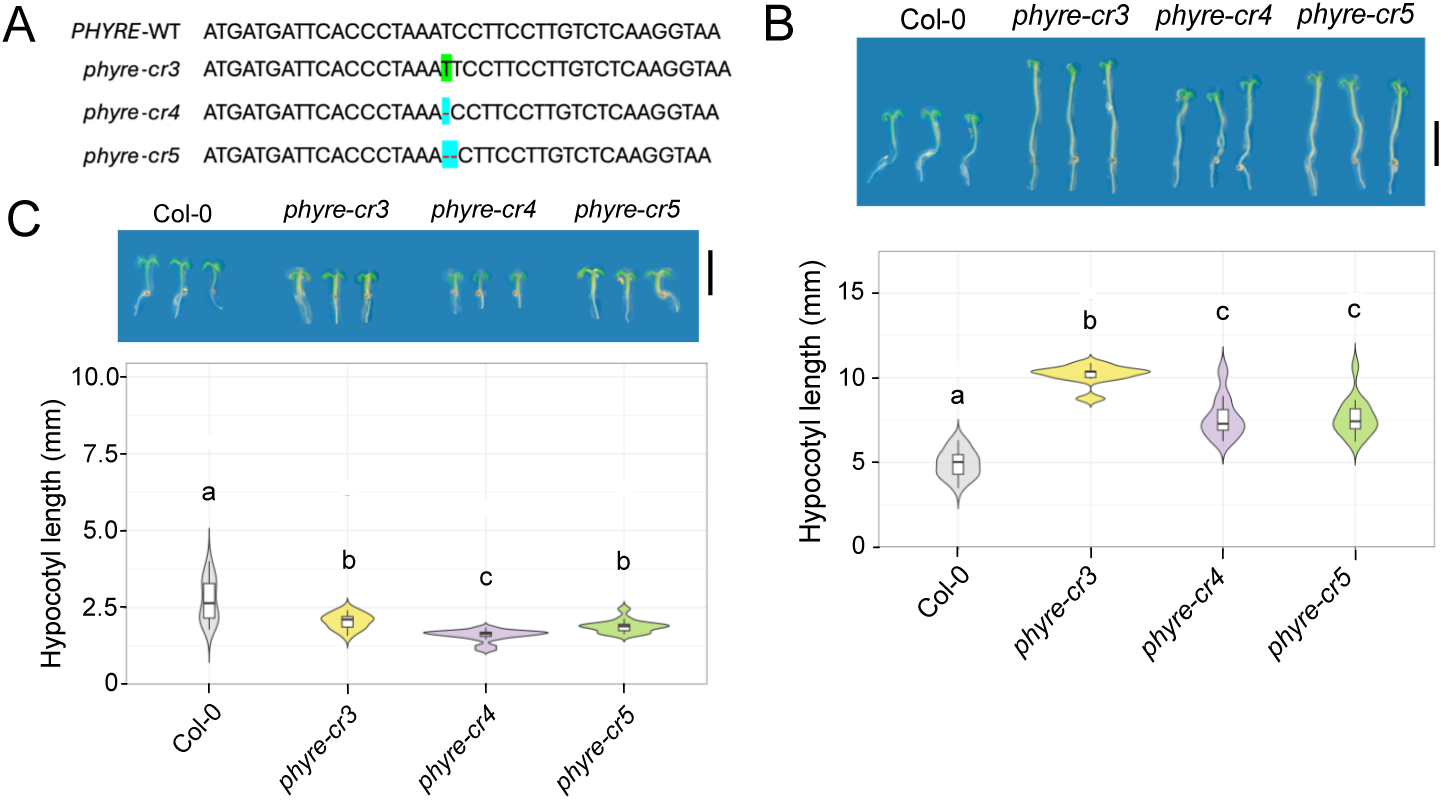
Hypocotyl phenotypes and phyB protein levels in *phyB-cr* and *phyre-cr* mutants. **(A)** Schematic showing the sequence information of the *PHYRE* uORF in the wild-type, *phyre-cr3*, *phyre-cr4*, and *phyre-cr5* mutants. The insertion in *phyre-cr3* is highlighted in green, and the deletions in *phyre-cr4* and *phyre-cr5* are indicated by blue highlighted red dashed lines. **(B-C)** Representative images and hypocotyl length quantification of 6-day-old seedlings of the indicated genotypes grown under **(B)** long-day white light or **(C)** long-day deep shade. Scale bars = 5 mm. Violin plots show the distribution of individual observations, with overlaid box plots indicating the median and interquartile range. Different letters indicate statistically significant differences among genotypes (adjusted P < 0.05), as determined using the Kruskal-Wallis test followed by Dunn’s post hoc test with Benjamini-Hochberg correction. n = 9-20 seedlings in **(B)** and 14-20 seedlings in **(C)**. **Note:** The representative image and violin plot data for Col-0 in **(B)** are reproduced from Figure 3D, as the genotypes in **(B)** were analyzed together with Col-0 as the reference control in the same experiment.

**Figure S3.**
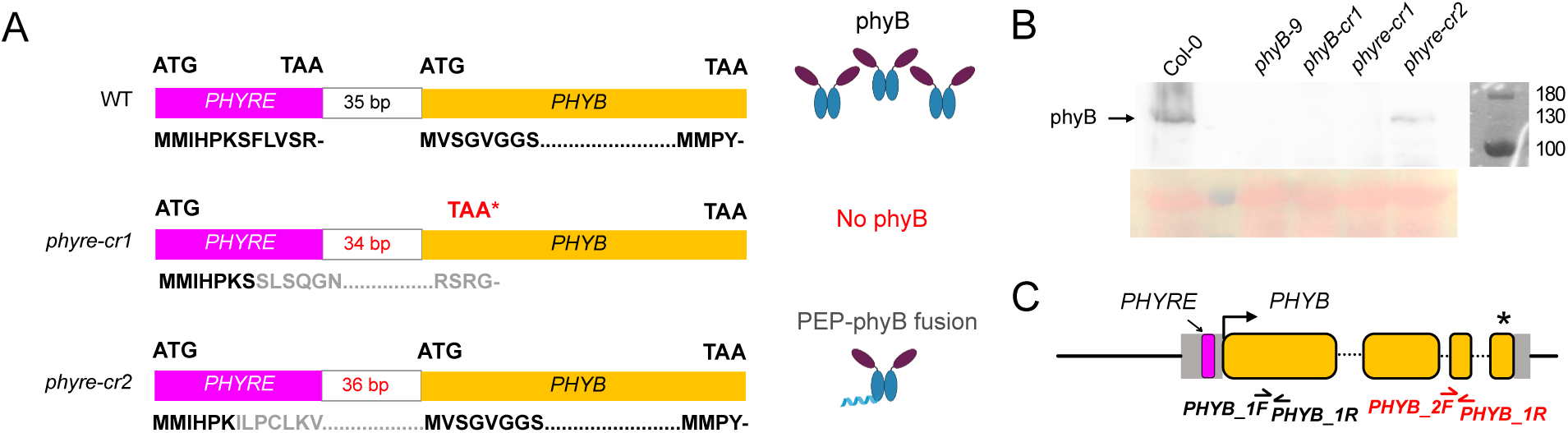
*PHYRE* and *PHYB* translation status in *phyre-cr* mutants. **(A)** Schematic showing translation status of *PHYRE* uORF and *PHYB* main ORF in wild type, *phyre-cr1* and *phyre-cr2*. Magenta and yellow rectangles represent the *PHYRE* and *PHYB* ORFs, respectively. Start and stop codons are indicated above the rectangles, and the corresponding amino acid sequences are shown below in black. Frameshift mutations in *phyre-cr1* and *phyre-cr2* eliminate the *PHYRE* stop codon and alter the spacing between the *PHYRE* stop codon and the downstream *PHYB* start codon. In *phyre-cr1*, the new reading frame extends into the *PHYB* coding sequence, resulting in loss of phyB protein due to the introduction of a premature stop codon (indicated in red). In *phyre-cr2*, the *PHYRE* and *PHYB* ORFs are brought into the same reading frame, allowing translation to extend into the downstream *PHYB* coding sequence and resulting in the production of an N-terminally extended phyB fusion protein. The altered amino acid sequences in *phyre-cr* mutants are shown in grey. **(B)** Western blot images showing phyB protein levels (129 kDa) in the total protein extracts from 7-day deep shade-grown seedlings of corresponding genotypes as indicated. Image of the corresponding ponceau stained blot (loading control) is shown below. **(C)** Schematic of the qPCR primer binding regions on the *PHYB* locus. The promoter and downstream regulatory regions are shown as black bars, 5’ and 3’ UTRs as grey boxes, and coding region as yellow boxes separated by introns (black dotted lines). The *PHYRE* uORF, *PHYB* start codon, and stop codon are indicated by a magenta rectangle, black arrowhead, and black asterisk, respectively.

**Figure S4.**
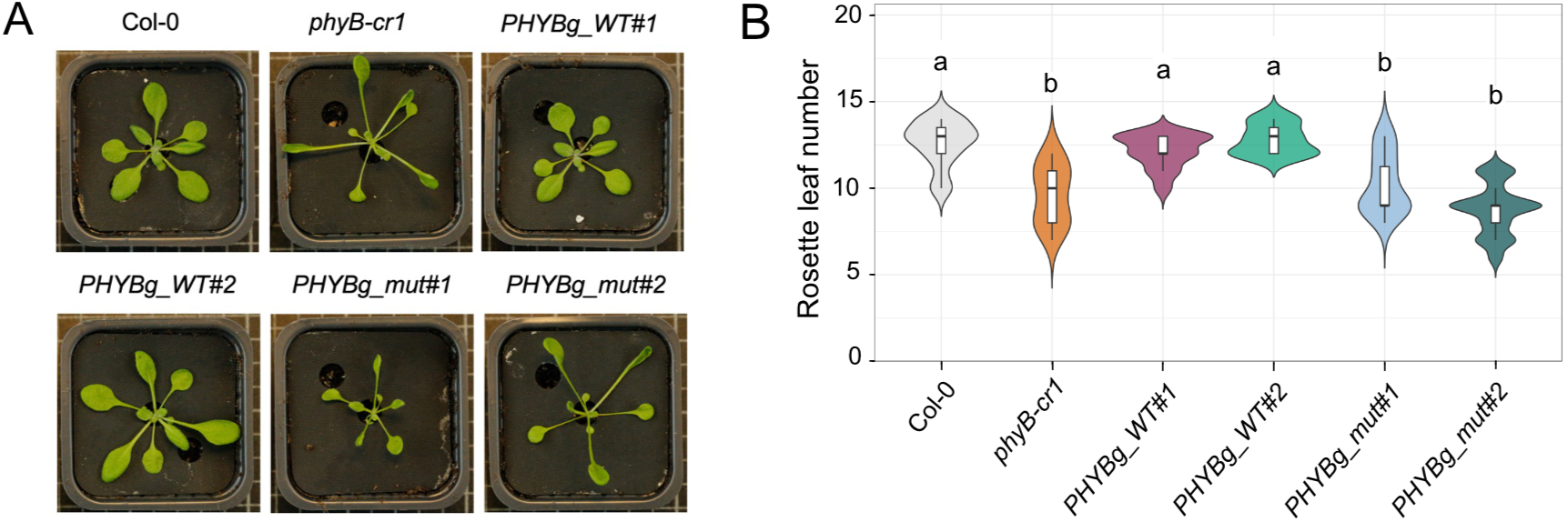
Flowering time phenotypes of *PHYB* complementation lines. **(A)** Representative rosette images of 21 day-old white light-grown plants of indicated genotypes. **(B)** Flowering time quantification indicated by the number of rosette leaves at bolting in the corresponding genotypes on the X-axis. Violin plots show the distribution of individual observations, with overlaid box plots indicating the median and interquartile range. Different letters indicate statistically significant differences among genotypes (adjusted P < 0.05), determined by Kruskal-Wallis test followed by Dunn’s post hoc test with Benjamini-Hochberg correction. n = 13-23 plants. **Note:** The representative image and violin plot data for Col-0 and *phyB-cr1* in **(A) and (B)** are reproduced from Figure 2F **and 2G** respectively, as the genotypes in **(B)** were analyzed together with Col-0 and *phyB-cr1* in the same experiment.

**Figure S5.**
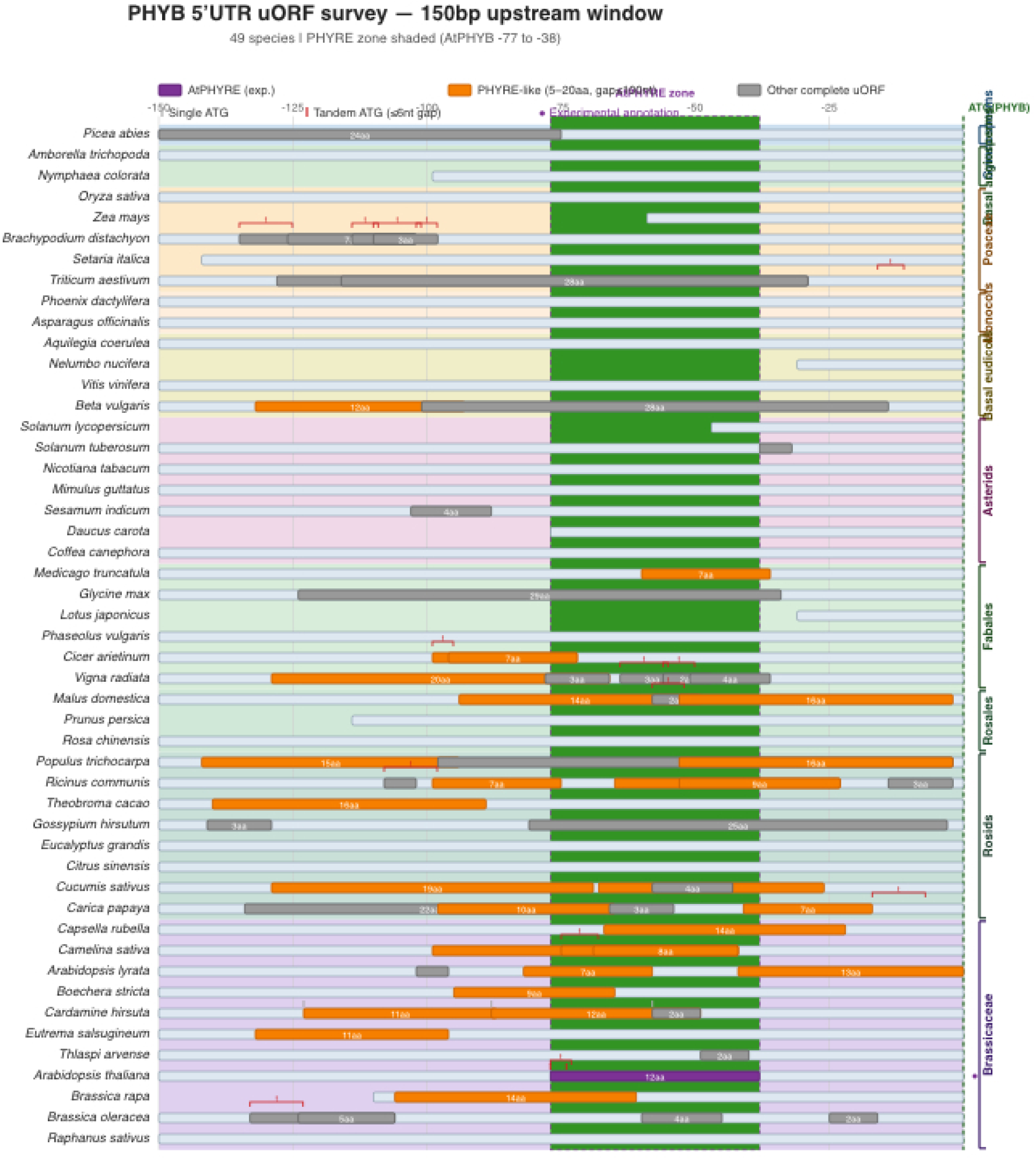
Comparative analysis of the *PHYB* 5′ untranslated regions across 49 land plant species. Genomic sequences spanning 150 nt upstream of the *PHYB* translation start site are shown as bars for 49 land plant species representing gymnosperms, monocots, basal eudicots, asterids, and rosids, with emphasis on *Brassicaceae*. Positions relative to the *PHYB* ATG are indicated above (0 to -150). The *Arabidopsis thaliana PHYRE* region (−74 to -36) is shaded green across all sequences to indicate the corresponding positional window. The experimentally validated *A. thaliana PHYRE* (39 bp) is highlighted in purple. *PHYRE-*like complete uORFs encoding 5-20 amino acids with a stop codon ≤100 nt upstream of the *PHYB* ATG are highlighted in orange, other complete uORFs encoding peptides shorter than 5 or longer than 20 amino acids are shown in grey.

## REFERENCES

1. Mahat, D.B., Salamanca, H.H., Duarte, F.M., Danko, C.G., and Lis, J.T. (2016). Mammalian Heat Shock Response and Mechanisms Underlying Its Genome-wide Transcriptional Regulation. Mol. Cell 62, 63–78. 10.1016/j.molcel.2016.02.025.

2. Schwanhäusser, B., Busse, D., Li, N., Dittmar, G., Schuchhardt, J., Wolf, J., Chen, W., and Selbach, M. (2011). Global quantification of mammalian gene expression control. Nature 473, 337–342. 10.1038/nature10098.

3. Liu, Y., Beyer, A., and Aebersold, R. (2016). On the Dependency of Cellular Protein Levels on mRNA Abundance. Cell 165, 535–550. 10.1016/j.cell.2016.03.014.

4. Ingolia, N.T., Ghaemmaghami, S., Newman, J.R.S., and Weissman, J.S. (2009). Genome-Wide Analysis in Vivo of Translation with Nucleotide Resolution Using Ribosome Profiling. Science (1979). 324, 218–223. 10.1126/science.1168978.

5. Ingolia, N.T. (2016). Ribosome Footprint Profiling of Translation throughout the Genome. Cell 165, 22–33. 10.1016/j.cell.2016.02.066.

6. Szavits-Nossan, J., and Ciandrini, L. (2020). Inferring efficiency of translation initiation and elongation from ribosome profiling. Nucleic Acids Res. 48, 9478– 9490. 10.1093/nar/gkaa678.

7. Pop, C., Rouskin, S., Ingolia, N.T., Han, L., Phizicky, E.M., Weissman, J.S., and Koller, D. (2014). Causal signals between codon bias, mRNA structure, and the efficiency of translation and elongation. Mol. Syst. Biol. 10, MSB145524. 10.15252/msb.20145524.

8. Juntawong, P., Girke, T., Bazin, J., and Bailey-Serres, J. (2014). Translational dynamics revealed by genome-wide profiling of ribosome footprints in *Arabidopsis*. Proceedings of the National Academy of Sciences 111. 10.1073/pnas.1317811111.

9. Liu, M., Wu, S., Chen, H., and Wu, S. (2012). Widespread translational control contributes to the regulation of Arabidopsis photomorphogenesis. Mol. Syst. Biol. 8, MSB201197. 10.1038/msb.2011.97.

10. Hinnebusch, A.G., Ivanov, I.P., and Sonenberg, N. (2016). Translational control by 5′-untranslated regions of eukaryotic mRNAs. Science (1979). 352, 1413–1416. 10.1126/science.aad9868.

11. Calvo, S.E., Pagliarini, D.J., and Mootha, V.K. (2009). Upstream open reading frames cause widespread reduction of protein expression and are polymorphic among humans. Proceedings of the National Academy of Sciences 106, 7507– 7512. 10.1073/pnas.0810916106.

12. von Arnim, A.G., Jia, Q., and Vaughn, J.N. (2014). Regulation of plant translation by upstream open reading frames. Plant Science 214, 1–12. 10.1016/j.plantsci.2013.09.006.

13. Sundaram, A., and Grant, C.M. (2014). A single inhibitory upstream open reading frame (uORF) is sufficient to regulate *Candida albicans GCN4* translation in response to amino acid starvation conditions. RNA 20, 559–567. 10.1261/rna.042267.113.

14. Gaba, A., Wang, Z., Krishnamoorthy, T., Hinnebusch, A.G., and Sachs, M.S. (2001). Physical evidence for distinct mechanisms of translational control by upstream open reading frames. EMBO J. 20, 6453–6463. 10.1093/emboj/20.22.6453.

15. Lin, Y., May, G.E., Kready, H., Nazzaro, L., Mao, M., Spealman, P., Creeger, Y., and McManus, C.J. (2019). Impacts of uORF codon identity and position on translation regulation. Nucleic Acids Res. 47, 9358–9367. 10.1093/nar/gkz681.

16. Dever, T.E., Ivanov, I.P., and Sachs, M.S. (2020). Conserved Upstream Open Reading Frame Nascent Peptides That Control Translation. Annu. Rev. Genet. 54, 237–264. 10.1146/annurev-genet-112618-043822.

17. Hendricks, S.B. (1960). Rates of Change of Phytochrome as an Essential Factor Determining Photoperiodism in Plants. Cold Spring Harb. Symp. Quant. Biol. 25, 245–248. 10.1101/SQB.1960.025.01.025.

18. Butler, W.L., Norris, K.H., Siegelman, H.W., and Hendricks, S.B. (1959). DETECTION, ASSAY, AND PRELIMINARY PURIFICATION OF THE PIGMENT CONTROLLING PHOTORESPONSIVE DEVELOPMENT OF PLANTS. Proceedings of the National Academy of Sciences 45, 1703–1708. 10.1073/pnas.45.12.1703.

19. Robson, PRH., Whitelam, G.C., and Smith, H. (1993). Selected Components of the Shade-Avoidance Syndrome Are Displayed in a Normal Manner in Mutants of Arabidopsis thaliana and Brassica rapa Deficient in Phytochrome B. Plant Physiol. 102, 1179–1184. 10.1104/pp.102.4.1179.

20. Sharrock, R.A., and Quail, P.H. (1989). Novel phytochrome sequences in Arabidopsis thaliana: structure, evolution, and differential expression of a plant regulatory photoreceptor family. Genes Dev. 3, 1745–1757. 10.1101/gad.3.11.1745.

21. Ni, W., Xu, S.-L., Tepperman, J.M., Stanley, D.J., Maltby, D.A., Gross, J.D., Burlingame, A.L., Wang, Z.-Y., and Quail, P.H. (2014). A mutually assured destruction mechanism attenuates light signaling in *Arabidopsis*. Science (1979). 344, 1160–1164. 10.1126/science.1250778.

22. Kircher, S., Kozma-Bognar, L., Kim, L., Adam, E., Harter, K., Schäfer, E., and Nagy, F. (1999). Light Quality–Dependent Nuclear Import of the Plant Photoreceptors Phytochrome A and B. Plant Cell 11, 1445–1456. 10.1105/tpc.11.8.1445.

23. Martínez-García, J.F., Huq, E., and Quail, P.H. (2000). Direct Targeting of Light Signals to a Promoter Element-Bound Transcription Factor. Science (1979). 288, 859–863. 10.1126/science.288.5467.859.

24. Hsu, P.Y., Calviello, L., Wu, H.-Y.L., Li, F.-W., Rothfels, C.J., Ohler, U., and Benfey, P.N. (2016). Super-resolution ribosome profiling reveals unannotated translation events in *Arabidopsis*. Proceedings of the National Academy of Sciences 113. 10.1073/pnas.1614788113.

25. Wu, H.-Y.L., Ai, Q., Teixeira, R.T., Nguyen, P.H.T., Song, G., Montes, C., Elmore, J.M., Walley, J.W., and Hsu, P.Y. (2024). Improved super-resolution ribosome profiling reveals prevalent translation of upstream ORFs and small ORFs in *Arabidopsis*. Plant Cell 36, 510–539. 10.1093/plcell/koad290.

26. Ji, Z. (2018). RibORF: Identifying Genome-Wide Translated Open Reading Frames Using Ribosome Profiling. Curr. Protoc. Mol. Biol. 124. 10.1002/cpmb.67.

27. Stuttmann, J., Barthel, K., Martin, P., Ordon, J., Erickson, J.L., Herr, R., Ferik, F., Kretschmer, C., Berner, T., Keilwagen, J., et al. (2021). Highly efficient multiplex editing: one-shot generation of 8× *Nicotiana benthamiana* and 12× Arabidopsis mutants. The Plant Journal 106, 8–22. 10.1111/tpj.15197.

28. Reed, J.W., Nagatani, A., Elich, T.D., Fagan, M., and Chory, J. (1994). Phytochrome A and Phytochrome B Have Overlapping but Distinct Functions in Arabidopsis Development. Plant Physiol. 104, 1139–1149. 10.1104/pp.104.4.1139.

29. Martínez-García, J.F., Gallemí, M., Molina-Contreras, M.J., Llorente, B., Bevilaqua, M.R.R., and Quail, P.H. (2014). The Shade Avoidance Syndrome in Arabidopsis: The Antagonistic Role of Phytochrome A and B Differentiates Vegetation Proximity and Canopy Shade. PLoS One 9, e109275. 10.1371/journal.pone.0109275.

30. Shinomura, T., Nagatani, A., Chory, J., and Furuya, M. (1994). The Induction of Seed Germination in Arabidopsis thaliana Is Regulated Principally by Phytochrome B and Secondarily by Phytochrome A. Plant Physiol. 104, 363–371. 10.1104/pp.104.2.363.

31. Gu, D., Wang, Y., Zhao, M., Chen, H., Wu, S., Jin, X., Deng, L., Ji, R., Xu, J., Zheng, F., et al. (2025). Phytochrome B stabilizes the KNOX transcription factor BP/KNAT1 to promote light-initiated seed germination in Arabidopsis thaliana. Plant Commun. 6, 101517. 10.1016/j.xplc.2025.101517.

32. Brogna, S., and Wen, J. (2009). Nonsense-mediated mRNA decay (NMD) mechanisms. Nat. Struct. Mol. Biol. 16, 107–113. 10.1038/nsmb.1550.

33. Hwang, Y., and Quail, P.H. (2008). Phytochrome-Regulated PIL1 Derepression is Developmentally Modulated. Plant Cell Physiol. 49, 501–511. 10.1093/pcp/pcn024.

34. Wiese, A., Elzinga, N., Wobbes, B., and Smeekens, S. (2004). A Conserved Upstream Open Reading Frame Mediates Sucrose-Induced Repression of Translation[W]. Plant Cell 16, 1717–1729. 10.1105/tpc.019349.

35. Ribone, P.A., Capella, M., Arce, A.L., and Chan, R.L. (2017). A uORF Represses the Transcription Factor AtHB1 in Aerial Tissues to Avoid a Deleterious Phenotype. Plant Physiol. 175, 1238–1253. 10.1104/pp.17.01060.

36. Wang, J., Liu, J., and Guo, Z. (2024). Natural uORF variation in plants. Trends Plant Sci. 29, 290–302. 10.1016/j.tplants.2023.07.005.

37. Rahmani, F., Hummel, M., Schuurmans, J., Wiese-Klinkenberg, A., Smeekens, S., and Hanson, J. (2009). Sucrose Control of Translation Mediated by an Upstream Open Reading Frame-Encoded Peptide . Plant Physiol. 150, 1356–1367. 10.1104/pp.109.136036.

38. Guerrero-González, M. de la L., Ortega-Amaro, M.A., Juárez-Montiel, M., and Jiménez-Bremont, J.F. (2016). Arabidopsis Polyamine oxidase-2 uORF is required for downstream translational regulation. Plant Physiology and Biochemistry 108, 381–390. 10.1016/j.plaphy.2016.08.006.

39. Casal, J.J. (2013). Photoreceptor Signaling Networks in Plant Responses to Shade. Annu. Rev. Plant Biol. 64, 403–427. 10.1146/annurev-arplant-050312-120221.

40. Martinez-Garcia, J.F., and Rodriguez-Concepcion, M. (2023). Molecular mechanisms of shade tolerance in plants. New Phytologist 239, 1190–1202. 10.1111/nph.19047.

41. Jung, J.-H., Domijan, M., Klose, C., Biswas, S., Ezer, D., Gao, M., Khattak, A.K., Box, M.S., Charoensawan, V., Cortijo, S., et al. (2016). Phytochromes function as thermosensors in *Arabidopsis*. Science (1979). 354, 886–889. 10.1126/science.aaf6005.

42. Legris, M., Klose, C., Burgie, E.S., Rojas, C.C.R., Neme, M., Hiltbrunner, A., Wigge, P.A., Schäfer, E., Vierstra, R.D., and Casal, J.J. (2016). Phytochrome B integrates light and temperature signals in *Arabidopsis*. Science (1979). 354, 897–900. 10.1126/science.aaf5656.

43. Kim, R.J.A., Fan, D., He, J., Kim, K., Du, J., and Chen, M. (2024). Photobody formation spatially segregates two opposing phytochrome B signaling actions of PIF5 degradation and stabilization. Nat. Commun. 15, 3519. 10.1038/s41467-024-47790-8.

44. Reed, J.W., Nagpal, P., Poole, D.S., Furuya, M., and Chory, J. (1993). Mutations in the gene for the red/far-red light receptor phytochrome B alter cell elongation and physiological responses throughout Arabidopsis development. Plant Cell 5, 147–157. 10.1105/tpc.5.2.147.

45. Grützner, R., and Marillonnet, S. (2020). Generation of MoClo Standard Parts Using Golden Gate Cloning. In, pp. 107–123. 10.1007/978-1-0716-0908-8_7.

46. Lampropoulos, A., Sutikovic, Z., Wenzl, C., Maegele, I., Lohmann, J.U., and Forner, J. (2013). GreenGate - A Novel, Versatile, and Efficient Cloning System for Plant Transgenesis. PLoS One 8, e83043. 10.1371/journal.pone.0083043.

47. Clough, S.J., and Bent, A.F. (1998). Floral dip: A simplified method for Agrobacterium-mediated transformation of Arabidopsis thaliana. Plant Journal 16, 735–743. 10.1046/j.1365-313X.1998.00343.x.

48. Mahboubi, A., Delhomme, N., Häggström, S., and Hanson, J. (2021). Small-scale sequencing enables quality assessment of Ribo-Seq data: an example from Arabidopsis cell culture. Plant Methods 17, 92. 10.1186/s13007-021-00791-w.

49. Dobin, A., Davis, C.A., Schlesinger, F., Drenkow, J., Zaleski, C., Jha, S., Batut, P., Chaisson, M., and Gingeras, T.R. (2013). STAR: ultrafast universal RNA-seq aligner. Bioinformatics 29, 15–21. 10.1093/bioinformatics/bts635.

50. Kubota, A., and Tajima, A. (2025). Click-qPCR: A Simple Tool for Interactive qPCR Data Analysis. Bio. Protoc. 15. 10.21769/BioProtoc.5513.

